# Tracking of Lineage Mass via Quantitative Phase Imaging and Confinement in Low Refractive Index Microwells

**DOI:** 10.1101/2024.03.27.587085

**Authors:** Jingzhou Zhang, Justin Griffin, Koushik Roy, Alexander Hoffmann, Thomas A. Zangle

## Abstract

Measurements of cell lineages is central to a variety of fundamental biological questions, ranging from developmental to cancer biology. However, accurate lineage tracing requires nearly perfect cell tracking, which can be challenging due to cell motion during imaging. Here we demonstrate the integration of microfabrication, imaging, and image processing approaches to demonstrate a platform for cell lineage tracing. We use quantitative phase imaging (QPI) a label-free imaging approach that quantifies cell mass. This gives an additional parameter, cell mass, that can be used to improve tracking accuracy. We confine lineages within microwells fabricated to reduce cell adhesion to sidewalls made of a low refractive index polymer. This also allows the microwells themselves to serve as references for QPI, enabling measurement of cell mass even in confluent microwells. We demonstrate application of this approach to immortalized adherent and nonadherent cell lines as well as stimulated primary B cells cultured *ex vivo*. Overall, our approach enables lineage tracking, or measurement of lineage mass, in a platform that can be customized to varied cell types.

## Introduction

Constructing cell lineages has been critical for improving our understanding of a variety of fundamental biological problems, such as the origins of cellular diversity and the stability of cell phenotypes [1]. In developmental biology, lineage tracing has been an effective tool for understanding stem cell proliferation and differentiation and how this leads to tissue development [2, 3]. In cancer, lineage tracing can help understand the origins of cellular heterogeneity and how this heterogeneity leads to tumor progression [4]. In applications to immunology, lineage tracing of B lymphocytes reveals the origins of cell fate variations, which could help understand the pathology of B cell-mediated diseases [5]. Lineage tracing of model organisms, for example yeast [6] and *C. elegans* [7], has been used to study cell-cell variability in cell fate decision making and molecular mechanisms during embryonic development.

Direct observation with automated label-free cell lineage tracing methods usually faces challenges due to cell segmentation and tracking when cells divide, migrate, and change in morphology [8]. Thus, some classic applications of lineage tracing often use cells with uniform size and morphology, such as yeast and bacteria, and often confine these to reduce the impact of motility. For example, cell segmentation and tracking can be easily performed with yeast cells confined within microchambers to create a monolayer with limited migration [6], or with rod-shaped bacteria confined within microchannels with a width closely matched to the cell diameter [9, 10]. Additionally, some mammalian cells (RBL and RAW 264.7) have also been confined into a uniform cylindrical geometry inside microchannels to perform lineage tracing [11]. However, such confinement of mammalian cells may affect cell behavior, for example by inducing asymmetric divisions [12]. Microwells offer another approach to separate cells and limit their movement [13], allowing for high throughput single cell analysis [14]. For example, microwells have been previously applied to track lymphocytes to show the growth fate competition and dynamics of activation and differentiation [5, 15, 16].

The fidelity of lineage tracing can be improved using markers, and so many lineage tracing methods use either genomic or fluorescent markers. When tracing cells within complex organisms, genomic methods can be used to infer cell lineage. For example, cell lineages can be reconstructed by tracing Cas9-induced mutations in a genetic barcode [17] [18]. Whenever possible, such as in cases of monolayer cells and organism with a small number of cells, more straightforward lineage tracing approaches are based on direct observation [5-8]. Highlighting cells with fluorescent labels can improve tracking efficiency [19]. However, to trace a large number of lineages at the same time, the number of fluorescence colors is limited to avoid interferences from different lineages.

One label-free approach to monitoring cells is quantitative phase imaging (QPI) [20, 21]. QPI measures the phase shift of light passing through cells and uses the known relationship between refractive index and cell mass density to calculate cell mass [20]. In cancer immunotherapy, QPI provides an efficient way for monitoring T cell/cancer cell interactions by measurements of cell biomass [22]. QPI also provides a label-free method for screening cancer cell drug response [23, 24]

Our lineage tracing approach is based on the combination of QPI with confinement in microwells. QPI enables us to use cell mass as new tracking parameter to create a label-free lineage tracing algorithm. Having mass as an additional dimension results in a simpler and more robust algorithm compared to tracking based on cell locations only [8]. To confine cells in the field of view during the multi-day imaging, we fabricated micro wells with MY133, a polymer with a reflective index close to water, for artifact-free imaging with QPI [25, 26]. We then use QPI measurement of cell mass as a label-free cell marker that does not photobleach or cause cell toxicity. Overall, when imaging with multiple locations of the microwell array, we are able to trace hundreds of lineages up to 5 generations within a single experiment setup. Cell mass data from confined microwells can also be used to track the total mass of individual lineages in cases where cells become confluent. Overall, the combination of low refractive index materials for cell confinement with QPI enables studies of the mass of cell lineages over time.

## Results & Discussion

### Cell imaging and lineage tracing

To track the lineage of both adherent and nonadherent cells, our approach combines QPI with microwells to contain cell lineages (**Figure 1**). We fabricate microwell structures using MY-133, a polymer with a refractive index is close to that of cell culture media. This results in structures that are mostly invisible when placed in water, and prevents artifacts or phase unwrapping that can occur with materials of higher refractive index [25, 26], while remaining biocompatible (**Figure S1**). We seed the microwells by directly adding the suspended cell culture solution onto our microwell device (**Figure 1A**). We target a seeding density such that there is approximately one founder cell per well area, to allow tracking of individual lineages and collect QPI data for 3-5 d (**Figure 1B**). Cell mass can then be computed as [20]:

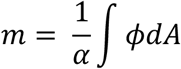

Where *m* is the cell dry mass, *α* is the specific refractive increment, which is a constant with a value approximately 1.8 × 10^−4^ m^3^/kg for most biological samples. *ϕ* is the measured phase shift in term of optical path length, and *A* is the area of the cell. Our image processing approach eliminates interference from the microwells and the background (**Figure 1C**). The mass of each founder cell and its descendants can then be measured by tracking cells contained within each microwell area with the image of the microwell itself removed automatically. For each well containing a founder cell, we track the founder cell and its descendants by QPI (**Figure 1D-F**). The mass growth profile of each individual cell within each lineage can then be computed by summing up pixels inside the individual cell area of the QPI-derived mass image (**Figure 1G**). These data show the expected doubling of cell mass with each generation as well as the increase in cell number due to proliferation. We then use the mass and location data from each cell to automatically link each pair of descendant sibling cells with the parent cells from which they originated. The result is a lineage tree for each tracked founder cell (**Figure 1H**).

**Figure 1:**
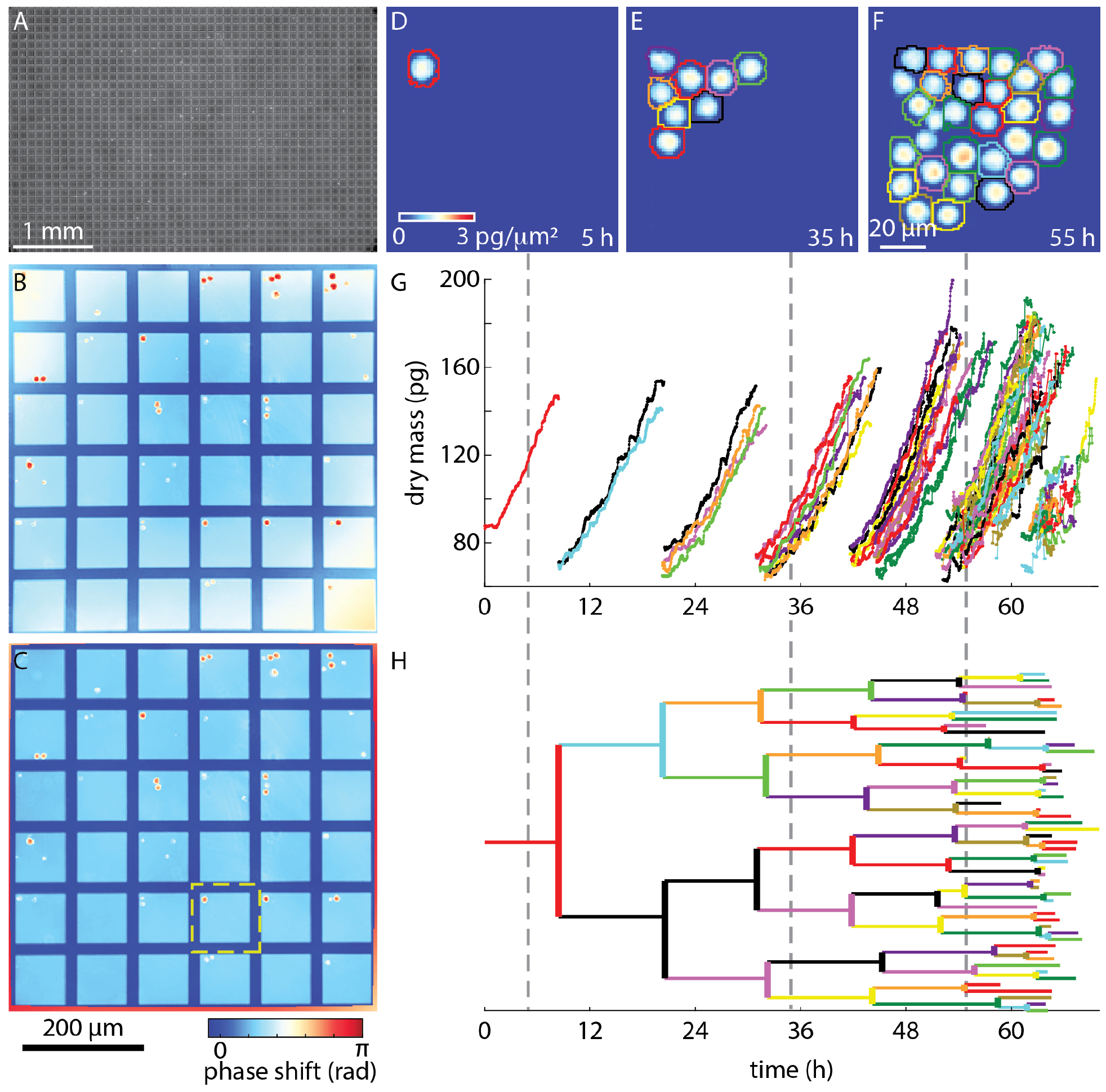
Long term lineage tracking with QPI and microwells. **A:** Microwells with cells seeded at the beginning of the experiment. Total area covered by microwells is 10 mm x 10 mm. **B:** Computed phase image of an individual field of view. **C:** Phase image of cells generated by performing background correction on B. Area inside the yellow dashed square is a single microwell. Colored outlines represent automated cell segmentation results. **D-F:** QPI of the single well from B after microwell subtraction showing a single cell and its lineage at 5, 35, and 55 hours. **G:** Cell mass over time and **H:** lineage tracking for 70 hours. Each colored track indicates the mass and lineage of a single cell. Colors in G-H correspond to cell segmentation labels in D-F at the times indicated by the gray dashed lines.

### Microwell fabrication

Our microwell fabrication and design process is specifically optimized for our image processing method to yield the maximum number of trackable cell lineages. A key feature of our microwell device is that the bottom of the microwells is directly exposed to the tissue culture treated dish substrate (**Figure 2**). This allows cells to adhere to the bottom of each microwell while preventing adhesion on the sidewalls. Preventing cells from growing or adhering on the sidewalls enables cell tracking with rapid, widefield imaging. To achieve microwell through holes, we used the capillary filling technique [27]. In contrast to adding MY 133 first, then pressing the mold into the material, the capillary filling method results in microwells without bottoms. This reduces any potential impact of MY 133 on cell growth and improves image quality by keeping cells within the same focal plane. We note, however, that with this method the MY 133 must be allowed to homogenize within the device overnight for adequate curing.

**Figure 2:**
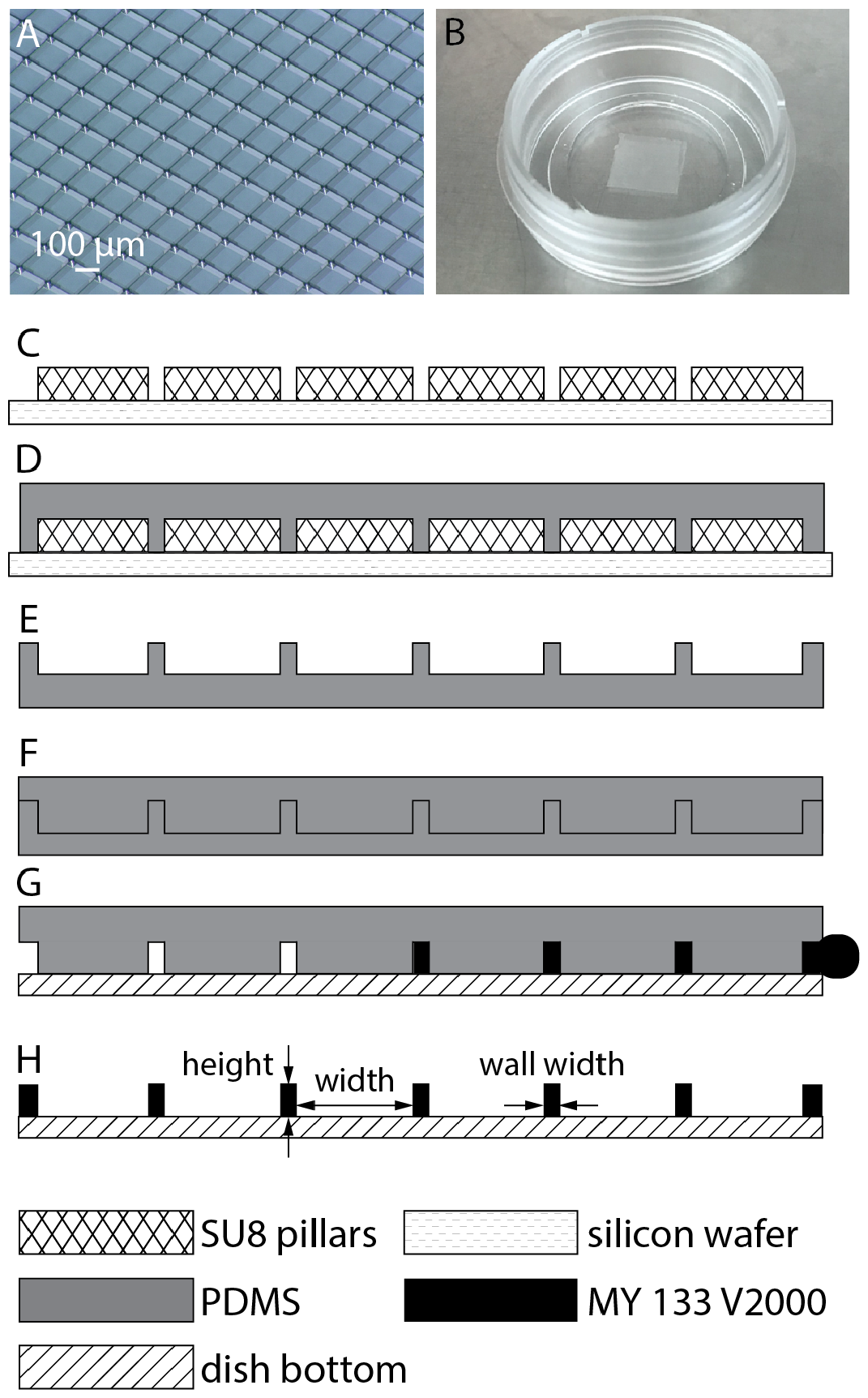
Fabrication process of microwell devices: **A:** 3 dimensional microscopy photo of our silicon wafer mold: SU-8 micro pillars build on silicon wafers by using photolithography. **B:** Photograph of finished MY 133 microwells device. The square area on the bottom of the dish is approximately 10 mm square and contains fabricated microwells. **C:** Schematic of SU8 micropillars built on a silicon wafer. Pillars are made instead of wells for durability of the SU8 mold. **D:** PDMS is poured and cured on the silicon wafer. **E:** Cured PDMS mold resulting from process D. **F:** The secondary PDMS mold is made by curing PDMS on the mold from D-E. This second mold will be pillars and used in a final molding step from MY133. **G:** The secondary mold from process F is placed in contact with the bottom of a 35 mm dish. MY-133 is added by capillary filling. **H:** The MY-133 is cured under UV light before lifting the secondary PDMS mold. The results are MY-133 microwells with exposed dish bottom for cell adhesion/growth.

The capillary filling method uses a PDMS stamp with a flat upper surface as the device mold. To make the stamp, we first fabricated SU8 micropillars on a silicon wafer using photolithography within our clean room facility (**Figure 2A, C**). The size of the micropillars will be the same size as the finished micro wells, which is designed to fit the desired number of microwells inside the field of view of our microscope (**Table 1**). These microwell sizes were chosen based on the typical cell area, to enable tracking of up to 5 generations (32 cells) within each microwell. After creating the PDMS mold (**Figure 2C-F**), we placed it on a tissue culture treated dish substrate and capillary fill the stamp with MY-133 by applying a droplet of uncured MY-133 on one side of the stamp. The uncured MY-133 spontaneously fills the stamp via capillary action (**Figure 2G**). After the MY-133 fills the stamp completely and has been allowed to homogenize, we cured the MY-133 with UV light and remove the PDMS stamp (**Figure 2F**). This results in the finished device of microwells inside a tissue culture treated dish (**Figure 2H**).

**Table 1.** Size of square microwells. Size is optimized to allow up to five generations of cells to be tracked. Depth of the wells is chosen to be high enough to prevent cells from escaping.

| Cell type | Width (μm) | Wall width (μm) | Height (μm) | Objective magnification | Number of wells in FOV | Imaging locations | Time per frame (min) | Imaging time (days) |
| --- | --- | --- | --- | --- | --- | --- | --- | --- |
| 70Z/3 | 140 | 20 | 45 | 40x | 9 | 40 | 6 | 4 |
| MCF7 | 155 | 20 | 40 | 60x | 1 | 18 | 10 | 5 |
| Mouse primary B cells | 85 | 20 | 45 | 20x | 36 | 19 | 2.5 | 5 |

### Image processing of microwell images

A key step in image processing of QPI data is the removal of background phase shifts present in raw phase images (**Figure 3A**). This step provides two functions: 1) it defines the location of ‘zero’ phase, from which other phase shifts can be measured, and 2) it corrects for any residual background phase shifts due to optical distortions or out of focus debris in the optical path. However, proper background correction requires identification of a zero phase reference region outside of the cell area that can account for any nonuniformities in the background phase shift. This can be challenging as cells become confluent. In our devices, cells preferentially adhere to the tissue culture-treated microwell bottoms rather than the fluorinated polymer sidewalls (**Figure 2**). As these microwells are fabricated using a low-refractive index polymer, we are able to avoid errors due to phase unwrapping at microwell edges [25, 28]. This means the repeating structure of the microwell walls is nearly cell free, giving us features that we leverage here for accurate QPI estimation of cell mass even as cells become confluent by aligning raw images to an average microwell image (**Figure S1**), then fitting a polynomial surface to microwell sidewalls (**Figure 3B**). We then subtract this phase shift (**Figure 3C**) from the QPI data for background correction and perform alignment to the microwell structures to correct for any variation in microwell positioning within the field of view from frame to frame (**Figure S2**). The result is a background corrected image of cells in microwells, from which we automatically detect wells with single founder cells for further analysis (**Figure 3D**).

**Figure 3.**
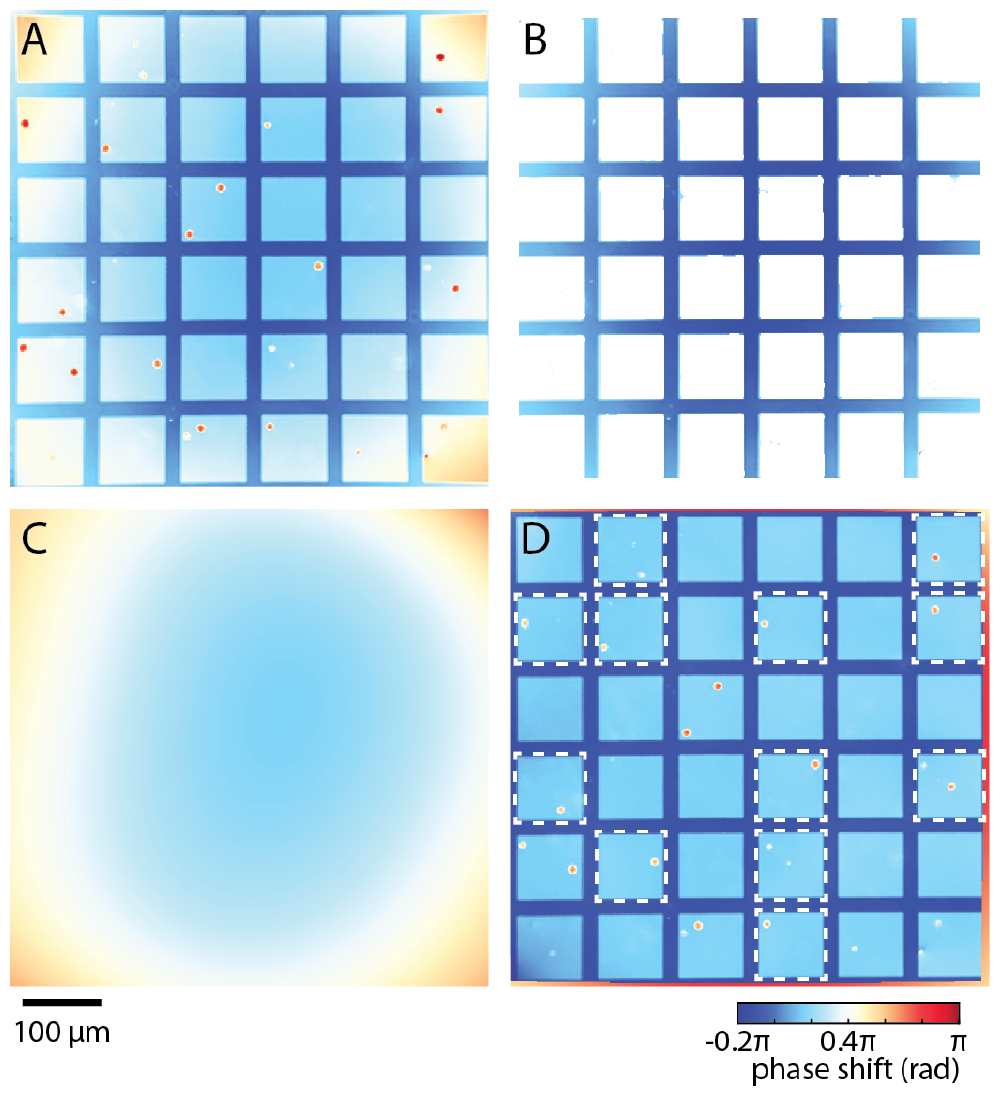
Image processing of microwell devices. **A:** Raw phase image showing microwell edges (straight line features), background phase shifts from the microscope system (notable as increased phase shift at the corners), and cells (small objects with large phase shift). **B:** Microwell edges are segmented from this image. **C:** A 4th order polynomial is fit to the edges of the microwells. **D:** When this polynomial surface is subtracted from the raw phase image, the background phase shifts are removed, resulting in flat bottom areas of each well. Wells with a single founder cell are identified from this image (outlined here with dashed white lines).

In addition to tracking non-adherent cells (**Figure 1**), QPI with low-refractive index microwells can also be used to track adherent cell lineages (**Figure 4**). In order to accommodate the typically larger spread area of adherent cells, we fabricated larger microwells for use with adherent cells (**Table 1, Figure 4A**). This allows tracking of cells from single founder cells (**Figure 4B**), through 8 and 16 cells in the lineage (**Figure 4C-D**). Mass of individual cells as well as cell lineages can be constructed from these data as well (**Figure 4E-F**).

**Figure 4.**
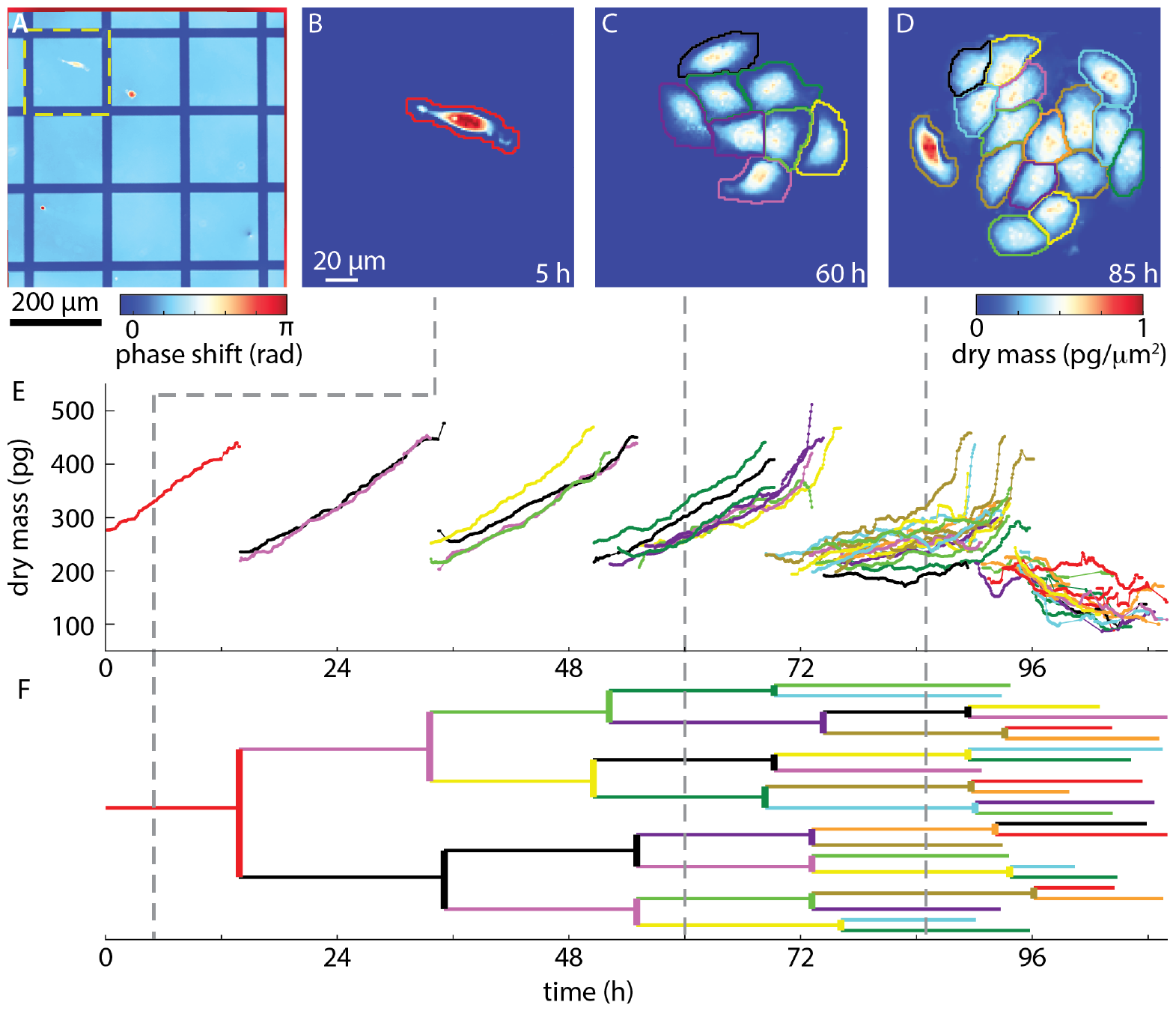
Tracking of adherent cell lineages with microwell confinement. **A:** Aligned and background corrected microwell image with one well containing a single founder MCF7 cell outlined (yellow dashed line). Colormap indicates phase shift due to both cells and microwells. **B-D:** QPI data of MCF7 cell lineage with automatic segmentation boundaries shown. Colormap of the base image shows cell mass distribution. **E:** Mass over time of cells tracked from lineage in the selected microwell. **F:** Cell lineage determined from cell tracking data. Colors in E-F correspond to cell segmentation labels in B-D at the times indicated by the gray dashed lines.

Though lineages can be constructed for single cells within our microwell devices for up to 5 generations (**Figures 1** and **4**), confinement of lineages within defined microwells also allows tracking of total lineage mass (**Figure 5**). This provides a robust measurement of lineage growth. In cases of robust growth and periodic cell doubling and division, lineage total mass shows a clear exponential increase of total mass over time (**Figure 5A,C**). When cells within a lineage start to die, tracking can be especially difficult (**Figure 5B**), even in sparse lineages (**Figure 5C**), due to the large shape changes and decreases in cell mass associated with cell death. In these cases, the lineage total mass can be used to effectively track lineages, without the need for precise tracking (**Figures 5D-F**). These results reveal distinct patterns of total mass accumulation for growing (**Figure 5D**), and dying (**Figure 5E-F**) lineages.

**Figure 5.**
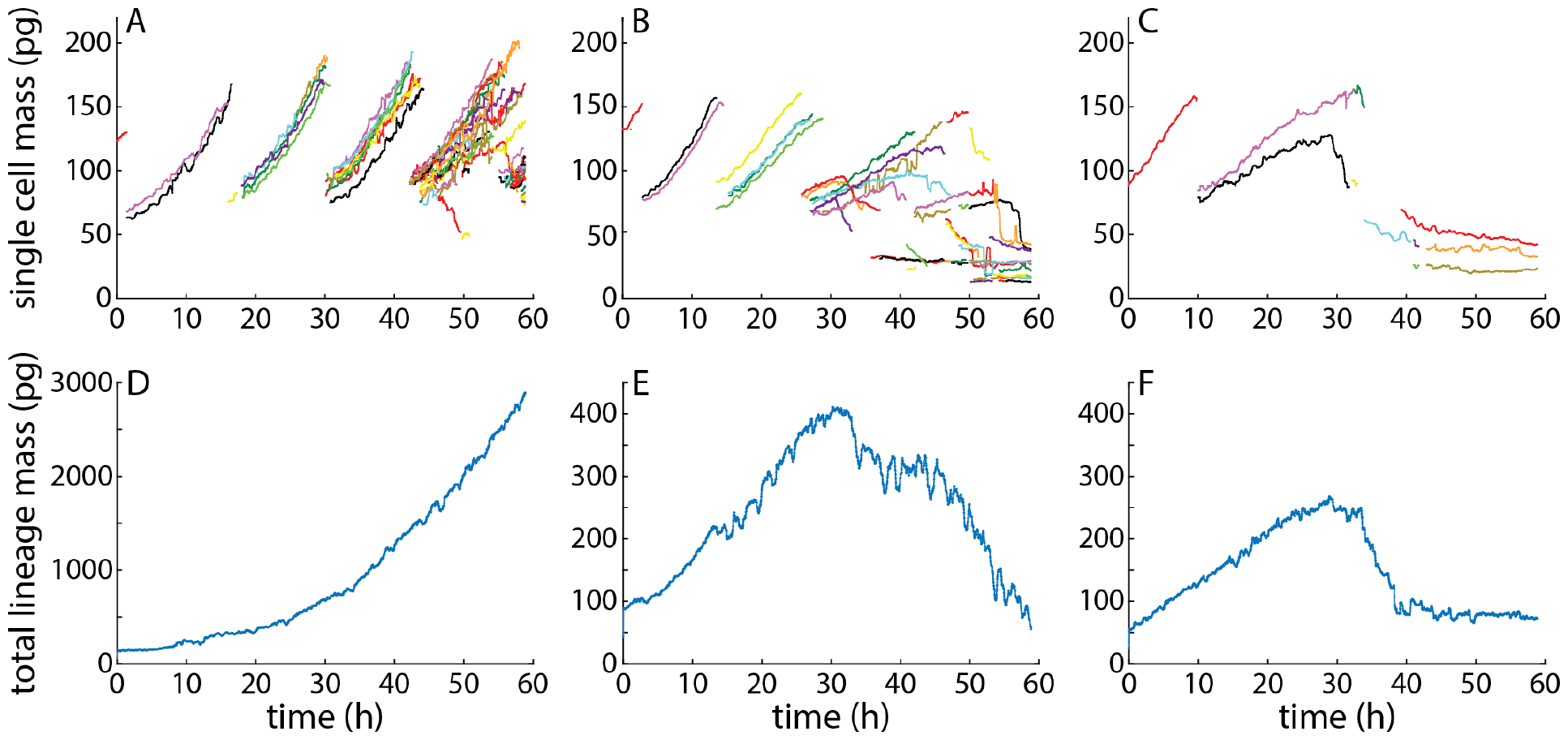
Measurement of total lineage mass using microwell confinement. **A:** Single cell mass over time for a growing lineage. **B:** Single cell mass over time for a gradually dying lineage. **C:** Single cell mass over time for a lineage that dies soon after the start of imaging. Each tracked cell is given a distinct color. **D-F:** Total mass of confined lineages corresponding to panels A-C. All lineages show 70Z/3 cells.

As a demonstration of the utility of our method to track both growth and death within lineages, we applied microwell-based tracking to mouse primary B cells stimulated *ex vivo* (**Figure 6**). B cells were CD43 depleted (**Figure S2**), plated in microwell devices (**Table 1**), and imaged via QPI for 120 h (**Figure 6A-D**). B cells imaged at maximal (**Figure 6B**) or near-maximal (**Figure 6D**) lineage mass show robust maintenance of cell size post division, as confirmed by single-cell tracking (**Figure 6E-F, Movies M1-M2**).

**Figure 6.**
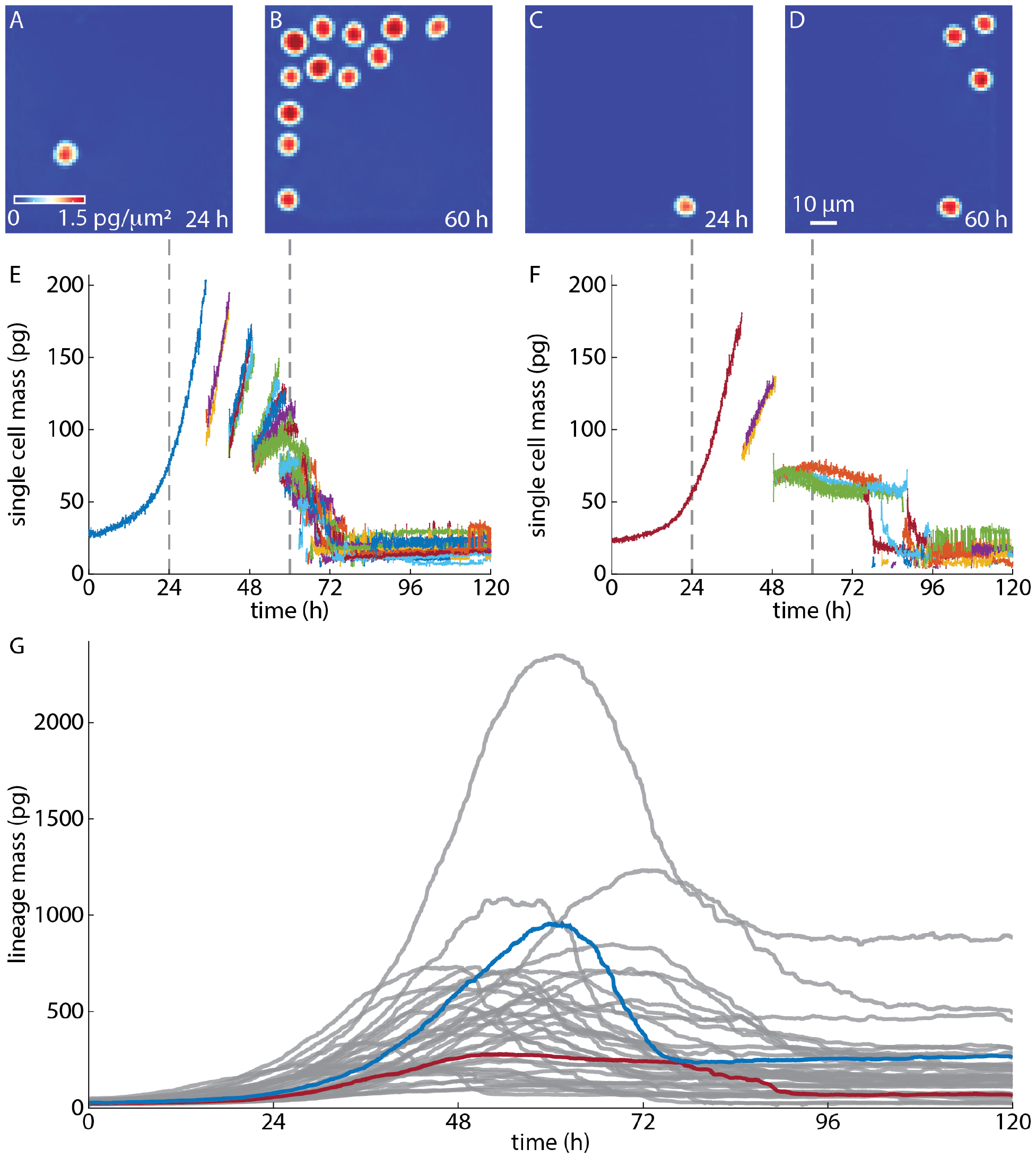
Primary B cell single cell and lineage mass. **A-D:** QPI data from two selected microwells showing CpG stimulated primary B cells and their progeny at 24 and 60 h after the start of imaging. **E:** Mass over time of tracked single cells from the microwell shown in **A-B. F:** Mass over time of the microwells shown in **C-D. G:** Total lineage mass over time for 35 lineages originating from single cells. Lineage from panel **E** is shown in blue. Lineage from panel **F** is shown in magenta.

However, during subsequent proliferation and death, single cell tracks can become fragmented, leading to errors in linage tracking (e.g. fragments at 80 h in **Figure 6F**). Therefore, total lineage mass provides a window into lineage growth dynamics that is unaffected by any errors in single-cell tracking (**Figure 6G**). Lineage total mass data reveal a maximum lineage mass of 5-fold or greater relative to initial mass and show a distinctive pattern of growth. These data may be used to study B cell growth and heterogeneity within individual lineages, which may lead to broader insights in immune system function [5].

## Conclusions

QPI is a powerful tool for monitoring the growth of live cells. In this study, we addressed the application of QPI for long term lineage tracking by combining QPI with a microfabricated device designed for and integrated into an image processing pipeline. With this method, we showed it is possible to automatically track the growth profile of adherent or nonadherent mammalian cell lineages for up to 5 generations, including from primary cells. Our approach leverages the cell mass data provided by QPI to provide another dimension for use in tracking. As a result, we capture cell mass data for each cell in constructed lineages, which might have the potential applications in the study of cell size control [10], and cell mass heredity [5] during proliferation.

Here, we utilize MY133 for fabrication of refractive index-matched microwells. We find that microwells made of MY 133 have three major advantages. First, the refractive index of MY 133 closely matches that of water, which avoids phase unwrapping issue when constructing the QPI image, even when cells are near the sidewalls. Second, the MY133 material is biocompatible, allowing cells to grow normally inside the wells. Third, the material has low adhesion, so cells adhere preferentially to the adhesive dish bottom, which is exposed by the through holes fabrication methodology. We also note that MY133 and QPI are both inherently compatible with fluorescence, potentially enabling study of lineages marked with fluorescent fusion proteins to study molecular regulation within cells [29].

Finally, we note that, in this study, we tailored the size of the microwells to balance throughput, in terms of number of lineages possibly measured, and lineage depth. In particular, we optimized the microwell size to fit up to 36 wells within the QPI field of view for application to nonadherent cells. However, this can still create issues with crowding as cell lineages expand. Therefore, larger wells could be used for deeper lineage tracking or longer term tracking with flow of media to feed cells, as required for more in depth studies of heritability of cell size. Alternatively, if shallower lineages are required, e.g. to track cells after one division, then smaller wells can be leveraged to increase throughput. In either case, this study highlights the applications possible when designing the whole process from device fabrication, through imaging, and image processing, to improve lineage tracing.

## Experimental

### Primary Mold Fabrication

We first designed microwell patterns using AutoCAD according to the device sizes listed in **Table 1**. SU-8 molds on silicon wafers were fabricated in the Utah NanoFab. We spin coated the silicon wafer with SU-8 to reach the desired average thickness.

### MY-133 microwell fabrication in cell culture Dish

PDMS (Sigma-Aldrich) at 10:1 (base: curing agent) was poured on the SU-8 micropillars and cured at 65 °C. We peeled PDMS microwells off the wafer, and then surface treated with 1H,1H,2H,2H-Perfluorooctyltrichlorosilane (Sigma-Aldrich) to serve as the mold for the PDMS micropillars stamp which was also treated with 1H,1H,2H,2H-Perfluorooctyltrichlorosilane (Sigma-Aldrich). The PDMS micropillars stamp was placed on the bottom of a 35 mm, polymer-bottom dish (ibidi), then one droplet (approximately 10 µL) of MY 133 V2000 (MY Polymers) was added to one edge of the stamp. Capillary flow then drew uncured MY 133 into the small gap between the stamp and dish bottom [1]. The filled stamps sat overnight to homogenize the components of MY 133, resulting in more even device curing. MY-133 was cured under 2 mL of water with an UV oven (Uvitron international Intelli-ray 400) for 1 h at 50% intensity and then an additional 1 h at 50% intensity after removing the PDMS stamp underwater. The final device was rinsed with ethanol.

### Cell Culture

We used 70Z/3 (ATCC#TIB-158) as a representative non-adherent cell line and MCF7 (ATCC# HTB-22) as a representative adherent cell line. Cells are first thawed and incubated at 37°C with 5% CO2 for 24 h. We used RPMI 1640 cell culture media with 10% FBS for 70Z/3 and Gibco DMEM with 10% FBS for MCF7.

### B cell isolation and culture

B cells were isolated from C57BL/6 mouse spleens using negative selection via CD43 (Ly-48) MicroBeads (Miltneyi Biotec, Gmbh, 130-049-801), LS Columns (Miltneyi Biotec, Gmbh, 130-042-401), and a MidiMACS™ Separator (Miltneyi Biotec, Gmbh) with isolation confirmed by flow cytometry for B220 (Rat monoclonal anti-mouse/human B220-FITC, BioLegend, Cat#103206) and CD43 (PE/Cy7 anti-mouse CD43 Antibody, BioLegend, Cat#143210). Cells were cultured using RPMI (Gibco, 11875-119) supplemented with 20 mM HEPES, penicillin-streptomycin (1X), 5 mM L-glutamine, β-ME (55 µM), and 10% FBS. B cells were stimulated by adding 250 ng/mL CpG (Invivogen, tlrl-1668) prior to imaging with QPI.

### Cell Plating

To reduce cytotoxicity, the finished microwell device was soaked in 200 mL of 100% ethanol for 2 d then transferred into 800 mL DI water and soaked for 3 d. To sterilize the device before plating we rinsed the microwells with 70% ethanol for about 10 seconds to remove air bubbles then submerged the dish and its lid under 70% ethanol solution before bringing it into a biosafety cabinet and rinsing 3x with autoclaved water and 2x with cell culture media. The target cell density was approximately one cell within the area of two wells. E.g. this translates to 34,000 70Z/3 cells and 12,000 MCF7 cells to cover the 750 mm^2^ ibidi dish bottom. After counting, cells were diluted to these target densities in 2 mL of cell culture media and added to the devices. 1-2% anti-clumping agent (Gibco) was added to 70Z/3 cells.

### Imaging

Locations with 2-15 wells within a field of view occupied by a single cell were selected for imaging resulting in 10-36 imaging locations per experiment. Images were acquired for 60 – 170 h using an Olympus IX83 microscope with on-stage incubator (OkoLab K301) at 37°C and 5% CO2. A Phasics SID4Bio-4MP camera (Phasics), 20x, 0.45 numerical aperture objective, and LED illumination at 623 nm (Thorlabs Solis) were used for imaging at 90 s (70Z/3), 300 s (MCF7), or 150 s (B cell) between frames. Raw interferograms were processed using Phasics Matlab SDF (Phasics) to compute phase images.

### Well alignment and background correction

One image with well-centered microwells and an empty top left corner was selected per microwell configuration. Images were corrected for lens distortion using the undistortImage function in Matlab (Mathworks). All other frames were aligned to this image using cross-correlation. Residual background phase shifts were removed by applying a 4th order polynomial fit to the microwell walls that was subtracted from the phase images. Microwells with a single cell in the first frame were automatically selected and segmented into individual image stacks for processing. Cells were then detected using Sobel edge detection and the background in each isolated microwell was then background corrected using a 4th order polynomial.

### Cell segmentation and tracking

Cells were detected by edge and magnitude threshold detection followed by segmentation using a watershed transform based on local maxima. To remove the resulting pixels between adjacent cells that were unassigned, images were scaled up by 2x and then the watershed mask was dilated by 1 pixel. This mask was then scaled down by 0.5x, resulting in a watershed mask with no gaps between adjacent cells. Mass was computed using an assumed specific refractive increment of 1.8 x 10^-3^ m^3^/kg [20, 30] before tracking using the Crocker and Grier algorithm [31] in 3-dimensions (x position, y position, mass). To improve track fidelity, the beginning and end of each track was found and connected using the Crocker and Grier algorithm [31]. This reconnection was run iteratively, with increases or decreases to the search radius depending on whether the tracking was successful (track matches were found) or unsuccessful (too many objects to count).

### Lineage tracking

Individual cell tracks were stored as cell objects containing cell properties, including heritage information. Cell lineages were constructed using a modification of the Crocker and Grier algorithm [31]. First, start and end points of cell tracks were found. The mass of cells at track end points was divided by 2 to account for splitting of cell mass at division. Cell end point data was also duplicated to allow for two daughters to connect to each potential parent. Then, these modified data arrays were tracked using the Crocker and Grier algorithm [31]. A division was detected, and cell object parameters were updated if two daughters were found for each parent. As with re-tracking, this process was run iteratively to improve lineage fidelity. Lineage trees were plotted using the Matlab-tree class [32].

### Total mass calculation

Edge and phase threshold detection followed by watershed segmentation was used to find cells within each background-corrected well. The mass of each detected cell within the wells was summed to produce a total mass per frame.

## Acknowledgements

This work was supported by the National Institutes of Health (R01AI132731 A.H.) and the University of Utah Office of the Vice President for Research (T.Z.). This work was performed in part at the Utah Nanofab Cleanroom sponsored by the John and Marcia Price College of Engineering College of Engineering and the Office of the Vice President for Research.

## Author Contributions

Conceptualization, formal analysis, validation, and writing—review and editing, software, data curation, visualization, and writing, J.Z. and T.A.Z.; methodology and investigation, J.Z, J.G.; interpretation of data, J.Z., T.A.Z., and K.R.; supervision, project administration, resources, and funding acquisition, A.H. and T.A.Z.; All authors have read and agreed to the final version of the manuscript.

## References

1. Papaioannou, V.E., Concepts of Cell Lineage in Mammalian Embryos. Curr Top Dev Biol, 2016. 117: p. 185–97.

2. Lu, R., et al., Tracking single hematopoietic stem cells in vivo using high-throughput sequencing in conjunction with viral genetic barcoding. Nature Biotechnology, 2011. 29(10): p. 928–933.

3. Kretzschmar, K. and Fiona, Lineage Tracing. Cell, 2012. 148(1-2): p. 33–45.

4. Navin, N.E. and J. Hicks, Tracing the tumor lineage. Mol Oncol, 2010. 4(3): p. 267–83.

5. Mitchell, S., et al., Nongenetic origins of cell-to-cell variability in B lymphocyte proliferation. Proceedings of the National Academy of Sciences, 2018. 115(12): p. E2888–E2897.

6. Falconnet, D., et al., High-throughput tracking of single yeast cells in a microfluidic imaging matrix. Lab Chip, 2011. 11(3): p. 466–473.

7. Bao, Z., et al., Automated cell lineage tracing in Caenorhabditis elegans. Proceedings of the National Academy of Sciences, 2006. 103(8): p. 2707–2712.

8. Al-Kofahi, O., et al., Automated cell lineage construction: a rapid method to analyze clonal development established with murine neural progenitor cells. Cell Cycle, 2006. 5(3): p. 327–35.

9. Baltekin, Ö., et al., Antibiotic susceptibility testing in less than 30 min using direct single-cell imaging. Proceedings of the National Academy of Sciences, 2017. 114(34): p. 9170–9175.

10. Si, F., et al., Mechanistic Origin of Cell-Size Control and Homeostasis in Bacteria. Current Biology, 2019. 29(11): p. 1760–1770.e7.

11. Varsano, G., Y. Wang, and M. Wu, Probing Mammalian Cell Size Homeostasis by Channel-Assisted Cell Reshaping. Cell Rep, 2017. 20(2): p. 397–410.

12. Lancaster, O.M., et al., Mitotic rounding alters cell geometry to ensure efficient bipolar spindle formation. Dev Cell, 2013. 25(3): p. 270–83.

13. Rettig, J.R. and A. Folch, Large-Scale Single-Cell Trapping And Imaging Using Microwell Arrays. Analytical Chemistry, 2015. 77(17): p. 5628–5634.

14. Zhang, J., et al., Microwell array chip-based single-cell analysis. Lab Chip, 2023. 23(5): p. 1066–1079.

15. Zhou, J.H.S., et al., Stochastically Timed Competition Between Division and Differentiation Fates Regulates the Transition From B Lymphoblast to Plasma Cell. Frontiers in Immunology, 2018. 9.

16. Zaretsky, I., et al., Monitoring the dynamics of primary T cell activation and differentiation using long term live cell imaging in microwell arrays. Lab on a Chip, 2012. 12(23): p. 5007.

17. Woodworth, M.B., K.M. Girskis, and C.A. Walsh, Building a lineage from single cells: genetic techniques for cell lineage tracking. Nature Reviews Genetics, 2017. 18(4): p. 230–244.

18. McKenna, A., et al., Whole-organism lineage tracing by combinatorial and cumulative genome editing. Science, 2016. 353(6298): p. aaf7907.

19. Hsu, Y.-C., Theory and Practice of Lineage Tracing. STEM CELLS, 2015. 33(11): p. 3197–3204.

20. Zangle, T.A. and M.A. Teitell, Live-cell mass profiling: an emerging approach in quantitative biophysics. Nature Methods, 2014. 11(12): p. 1221–1228.

21. Park, Y., C. Depeursinge, and G. Popescu, Quantitative phase imaging in biomedicine. Nature Photonics, 2018. 12(10): p. 578–589.

22. Zangle, T.A., et al., Quantifying biomass changes of single CD8+ T cells during antigen specific cytotoxicity. PLoS One, 2013. 8(7): p. e68916.

23. Reed, J., et al., Rapid, Massively Parallel Single-Cell Drug Response Measurements via Live Cell Interferometry. 2011. 101(5): p. 1025–1031.

24. Chun, J., et al., Rapidly quantifying drug sensitivity of dispersed and clumped breast cancer cells by mass profiling. Analyst, 2012. 137(23): p. 5495–8.

25. Kim, D.N.H., et al., Soft lithography fabrication of index-matched microfluidic devices for reducing artifacts in fluorescence and quantitative phase imaging. Microfluidics and Nanofluidics, 2017. 22(1): p. 1–11.

26. Polanco, E.R., N. Western, and T.A. Zangle, Fabrication of refractive-index-matched devices for biomedical microfluidics. Journal of Visualized Experiments, 2018. 139: p. e58296.

27. Masters, T., et al., Easy fabrication of thin membranes with through holes. Application to protein patterning. PLoS One, 2012. 7(8): p. e44261.

28. Polanco, E.R., J. Griffin, and T.A. Zangle, Fabrication and Bonding of Refractive Index Matched Microfluidics for Precise Measurements of Cell Mass. Polymers (Basel), 2021. 13(4).

29. Roy, K., et al., A Regulatory Circuit Controlling the Dynamics of NFkappaB cRel Transitions B Cells from Proliferation to Plasma Cell Differentiation. Immunity, 2019. 50(3): p. 616–628 e6.

30. Ross, K.F.A., Phase Contrast and Interference Microscopy for Cell Biologists. 1967, New York: St. Martin’s Press.

31. Crocker, J.C. and D.G. Grier, Methods of Digital Video Microscopy for Colloidal Studies. Journal of Colloid and Interface Science, 1996. 179: p. 298–310.

32. Tinevez, J.-Y., matlab-tree. 2014: https://github.com/tinevez/matlab-tree.

