## Supplementary material for "Tracking of Lineage Mass via Quantitative Phase Imaging and Confinement in Low Refractive Index Microwells": Supplental information

1. Department of Chemical Engineering, University of Utah; 2. Division of Microbiology and Immunology, Department of Pathology, School of Medicine, University of Utah; 3. Signaling Systems Laboratory, Institute for Quantitative and Computational Biosciences and Department of Microbiology, Immunology, and Molecular Genetics, University of California, Los Angeles, Los Angeles, CA; 4. Huntsman Cancer Institute, University of Utah

**Contents:**

Supplemental Figures 1-2
Legends for Movies M1-M2

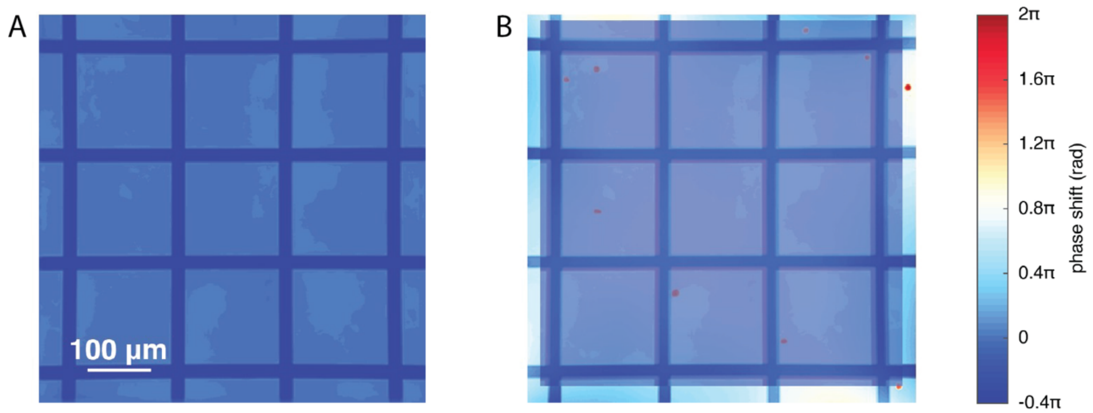


**Figure S1. Alignment of microwell images using repeating structure of microwells. A:** Empty microwell image generated from averaging regions without cells from collected QPI data. **B:** The central portion of the empty microwell image is then aligned to each frame containing QPI data. This results in an aligned subimage of each frame that is used for subsequent processing.

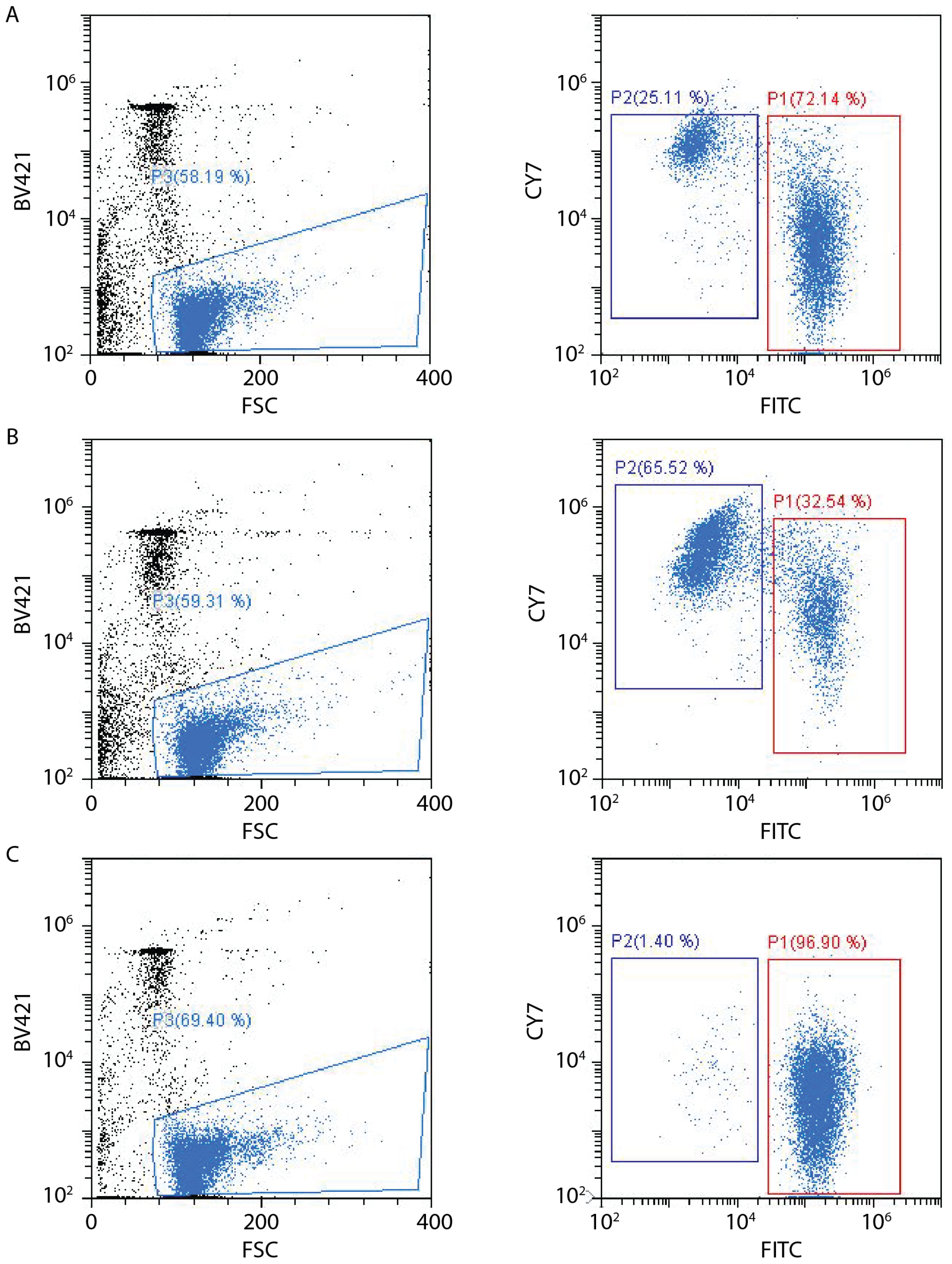


**Figure S2: Flow cytometry of sorted primary B cells isolated from mouse spleen. A:** Whole spleen isolate pre-sort. **B:** CD-43 positive sort of cells captured from the magnetic bead separation column. **C:** CD-43 negative sorted cells as effluent form the magnetic bead separation column. Left column shows DAPI (BV421) vs. forward scatter used to gate on live cells. Right column shows CD43 (Cy7 conjugated) vs. B220 (FITC conjugated) to identify B220+ CD43- subpopulation. Percentage of events falling within indicated gating shown on the individual plots.

**Movie M1:** QPI data of primary B cell lineage shown in Figure 6A-B over time. Red outlines show automatically generated segmentation, black points indicate cell centroids used for tracking, and numbers indicate automatically generated cell identification numbers used in tracking.

**Movie M2:** QPI data of primary B cell lineage shown in Figure 6C-D over time. Red outlines show automatically generated segmentation, black points indicate cell centroids used for tracking, and numbers indicate automatically generated cell identification numbers used in tracking.
